# BEstimate: a computational tool for the design and interpretation of CRISPR base editing experiments

**DOI:** 10.1101/2025.05.19.654892

**Authors:** Cansu Dinçer, Bo Fussing, Mathew J. Garnett, Matthew A. Coelho

## Abstract

CRISPR base editors enable scalable targeted DNA mutagenesis and are a powerful tool for the interpretation of gene function, the study of variants of unknown significance, and disease modelling. Existing guide RNA (gRNA) design tools focus on engineering a small number of targeted variants in a given gene or lack comprehensive functional annotation of target sequences. Here we developed BEstimate, a flexible computational pipeline that systematically specifies base editor gRNA target sites, generates gRNA off-target predictions, and rich functional, structural and clinical annotations of targeted edits. BEstimate accommodates custom gRNA design against variant alleles, improving editing accuracy and facilitating the programmed reversion of disease variants. BEstimate is a freely available, versatile tool for designing gRNA libraries and analysing base editor screens.

## Background

Genome editing has the potential to inform gene and gene variant function at scale (1,2) with implications for our understanding and treatment of genetic diseases (3–6). The continuous development in genome editing through Clustered Regularly Interspaced Short Palindromic Repeats (CRISPR) tools has rapidly expanded the capabilities of genome engineering. Specifically, base editing enables programmable single-nucleotide modifications without introducing double-strand breaks (7–10). This technology offers broad applications in both basic and translational research, such as functional genomics and therapeutic development. Examples of base editing applications in functional genomics include the study of variant effects in primary human T cells on their cytotoxic functionality and to pinpoint mutations that tune immune responses (11). Similarly, *JAK1* mutagenesis with base editing predicted variants altering IFN-γ pathway activity in colorectal cancer cells (12). Additionally, mutagenesis of 11 genes across four cancer cell lines with base editors elucidated variants related to sensitivity or resistance to cancer therapies and generated a systematic functional map of alterations (13). Base editing screening on *WRN* indicated the helicase domain as the primary therapeutic target in microsatellite-unstable (MSI) cancer cell lines (14). For therapeutic purposes, base editors have been successfully used to correct disease-causing mutations in sickle cell disease, β-thalassemia and Alpha-1 Antitrypsin Deficiency (15–17).

Base editors comprise a catalytically impaired Cas9 (typically Cas9 nickase or dead Cas9/dCas9) and a nucleotide deaminase. Using a dCas9 provides recognition of the targeted sequence via gRNAs without causing double-strand breaks, which can be deleterious in some cells. By overcoming the requirement for Homology-Directed Repair (HDR)-dependency and insertion and deletion (indel) formation, base editors can generate mutations on the targeted site through deamination (18). With dCas9 and deaminase, base editors facilitate the precise installation of single-nucleotide variants (SNV) within a defined editing window adjacent to the protospacer directed by gRNAs. There are several types of base editors with different specificities, enabling a range of edits to be introduced for each nucleotide position. Cytosine Base Editors (CBEs) use cytosine deaminase fused to Cas9 nickase leading to conversion from cytosine to thymine (C>T) (10,19); Adenine Base Editors (ABEs) use adenine deaminase leading to an adenine to guanine (A>G) conversion (9), and lastly Glycosylase Base Editors (CGBEs) employ both cytosine deaminase and uracil-DNA glycosylase for cytosine to guanine (C>G) transversions (20,21).

Base editors offer scalable, affordable, multiplexed genome editing in a range of cell types (22–24). Given their broad applications in generating and reverting variants, a robust tool for designing and annotating base editor gRNA libraries is essential. Base editors cannot target all positions on the genome due to the limitation of PAM preference (9,10), activity window restriction, and the nature of the deamination reaction (25). Also, gRNAs can result in off-target and bystander mutations, which should be carefully considered in experimental design and gRNA selection (26,27). Technological improvements have led to base editors with less stringent PAM sequences, reduced off-targets, more precise editing windows, and different nucleotide conversions, such as cytosine to guanine (28–33). Moreover, information about the potential functional consequences of the generated alterations allows experiment-specific library design and *in silico* analysis of the sequence of interest. In addition to selecting favourable gRNAs with minimal off-target effects, it can be useful to prioritise programmed DNA variants according to their potential functional, clinical and structural consequences. It is, therefore, vital to have a robust and flexible tool that provides customisation options for Cas9 and base editor enzymes, off-target information, and annotation of mutational consequences.

Current tools can identify either gRNAs targeting given DNA sequences (34–36) or gRNAs which can target provided alterations (34–37) **(Additional file 1: Table S1)**. While some tools have limited flexibility on nuclease, PAM and activity window selection (38), most tools only accommodate currently available base editors, such as SpCas9 NGG. Moreover, they lack a comprehensive annotation of gRNA effects. Thus, we developed BEstimate, a Python package that systematically analyses gRNA target sites across reference or mutated sequences for fully customised base editors, PAM, and activity window, and comprehensively annotates functional, structural and clinical consequences of programmed edits and provides information on gRNA off-targets. BEstimate provides *in silico* analysis of the sequences to identify positions that can be editable or alterations that can be correctable by base editors, as well as their features before starting experiments. Therefore, because of its flexible design and detailed annotations, it provides information ranging from gene sequence to clinical consequences of predicted edits and is readily extendable to accommodate future improvements in base editing enzymes.

## Results

We have previously used a prototype version of BEstimate to design three independent gRNA libraries targeting 20 genes (12–14). Here, BEstimate is implemented as a Python package retrieving real-time up-to-date information with *Ensembl* Application Programming Interface (API) (39,40) and *Uniprot* Representational State Transfer (REST) API (41,42) to collect and manipulate gene sequences, identify gRNA sequences for fully customisable base editors and comprehensively annotate them with functional, structural and clinical consequences **(Fig. 1)**. Here we demonstrate the use of BEstimate through three case studies, each highlighting different functions of the tool.

**Fig 1.**
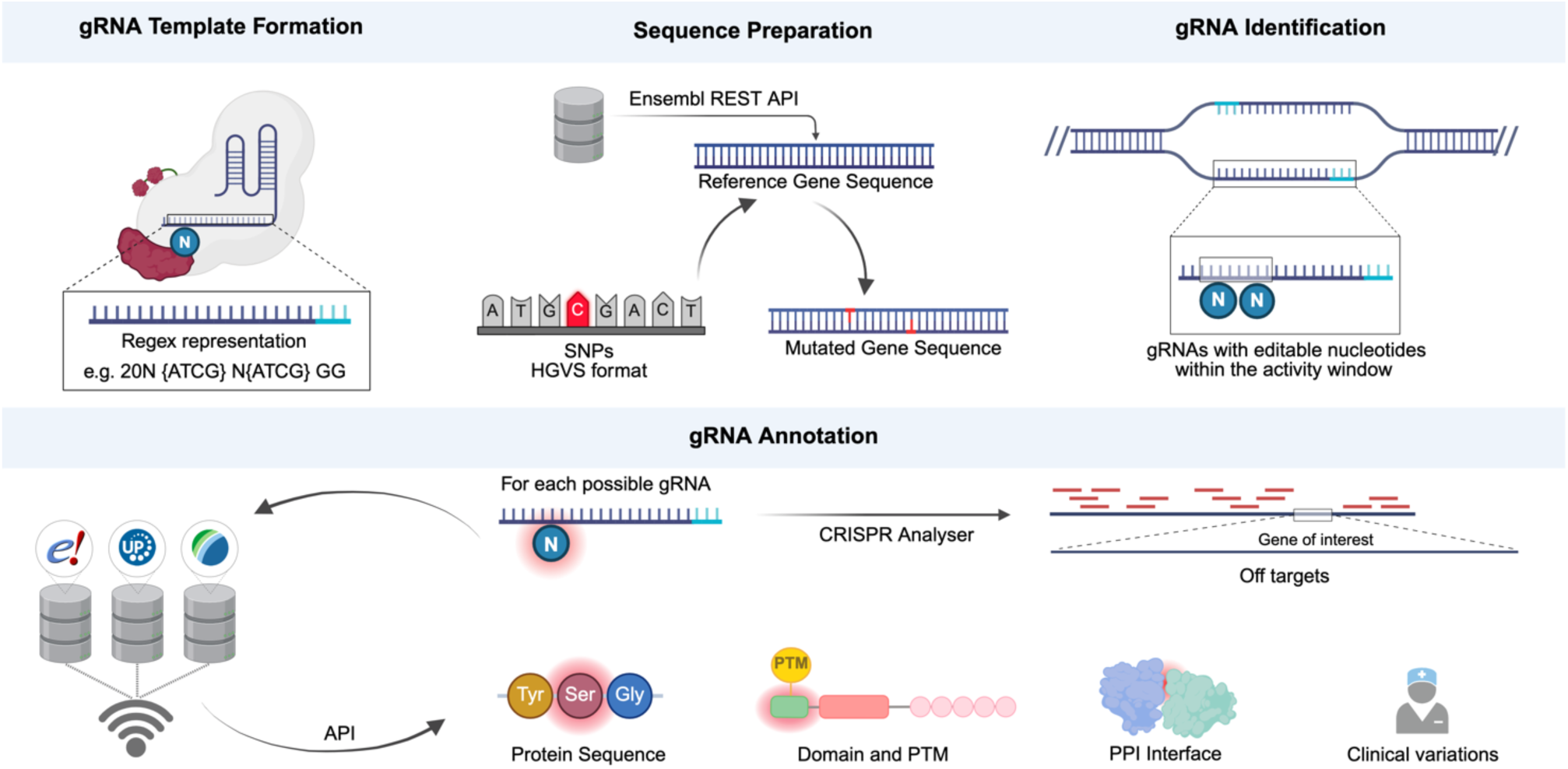
BEstimate is a flexible tool for gRNA design and annotation of edits. Given user-defined input, BEstimate (i) designs a regular expression (regex) representation of the gRNA, (ii) searches sites across the reference or altered gene sequence, aligning with the gRNA regex representation, (iii) finds genomic coordinates of gRNA and editable sites, (iv) annotates functional, structural and clinical consequences of potential gRNAs and gRNA off-target scores across the reference genome. Created in bioRender.

### Case study I: Designing base editor libraries with reference and model-specific gene sequences

A key application for base-editing is the analysis of variants involved in cancer and cancer drug resistance (13). To facilitate this and to demonstrate the utility of BEstimate, and extending previous gRNA libraries from BEstimate (12–14), we designed a cytosine and adenine NGG- and NG PAM-specific base editors gRNA library targeting 50 frequently mutated cancer genes derived from the *cBioPortal* pan-cancer study **(Additional files 2 and 3)** (43,44). To highlight the base editor coverage across nucleotide and amino acid sequences, we leveraged BEstimate for *in silico* analysis, collecting editable sites and consequences of the generated alterations on the genome and translated proteins. For the selected 50 genes with NGG-PAM and 3-9 activity window, we covered an average of 13.14% of nucleotide positions (standard deviation (σ) 5.76%) for cytosine, and 15.39% of nucleotide positions (σ=2.01%) for adenine base editors **(Fig. 2a)**. Leveraging a less stringent NG-PAM, there was an increase in the editable nucleotide coverages to 34.08% (σ=8.22%) for cytosine, and 42.53% (σ=3.20%) for adenine base editors **(Fig. 2b)**. At the amino acid level, we observed a coverage of the NGG-PAM library of 29.36% (σ=10.12%) for cytosine, and 29.10% (σ=6.03%) for adenine base editors **(Fig. 2a)**. Similar to the nucleotide coverage, NG-PAM specific base editors increased the amino acid coverage of cytosine and adenine base editors to 64.66% (σ=8.58%) and 66.09% (σ=4.10%), respectively **(Fig. 2b)**. Across all editable amino acids, 11.24% (σ=3.98%) and 38.99 (σ=3.75%) of them were shared between both base editors with NGG- and NG-PAM specificities, respectively **(Fig. 2ab)**. Using both base editors combined, the predicted average edited amino acid coverage per gene increased to 47.22% for NGG and 91.76% for NG PAM-specific base editors with a 3-9 activity window. These results demonstrate the value of using multiple base editors with permissive PAM sequences to increase the coverage of amino acid positions and achieve an overall high level of saturation.

**Fig 2.**
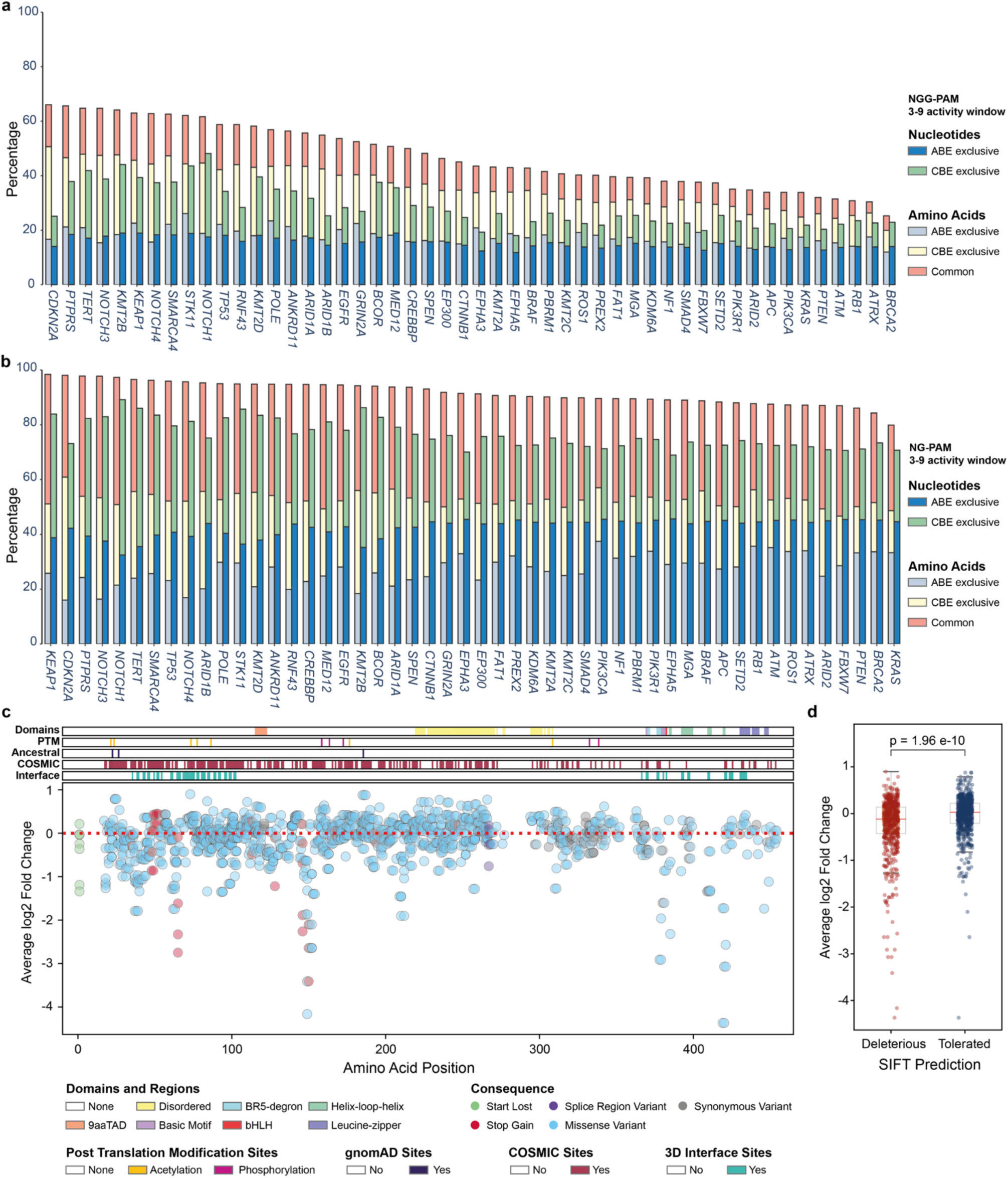
Representations of gRNA design and *in silico* annotation through BEstimate. **(a and b)** Barplot of **(a)** NGG-PAM specific cytosine (CBE) and adenosine (ABE) base editor coverage and **(b)** NG-PAM specific cytosine (CBE) and adenosine (ABE) base editor coverage on nucleotide and amino acid levels across 50 cancer genes. When an amino acid can be targeted only with one base editor, it was assigned as exclusive to the corresponding base editor; otherwise, it was labelled as commonly targeted. **(c)** Scatterplot of average log2 fold change (LFC) of MYC base editor screens with functional annotation of the alterations. The colour of dots represents the predicted mutational consequence of the gRNA. The top annotations indicate the domains and regions, post-translational modification sites of the corresponding amino acid positions, and whether these positions are in the Genome Aggregation Database (gnomAD), Catalogue of Somatic Mutations in Cancer (COSMIC), or on the protein-protein interaction (PPI) sites from Interactome Insider. **(d)** Average log2 fold change comparison between gRNAs predicted as deleterious and tolerated by the Sorting Intolerant From Tolerant (SIFT) algorithm. The boxplot represents the values between the 25th and 75th quantiles, where the red line indicates the median value. The p-value indicates the significance based on the two-way Mann-Whitney U Test.

Existing variants in target genes may negatively impact gRNA efficiency or create *de novo* PAM sequences. We designed BEstimate to generate gRNA sequences that account for SNVs within genes, including disease-associated variants. For example, we designed an NG PAM-specific ABE gRNA library for the *PIK3CA* gene specific to the head and neck cancer cell line, CAL-33 **(Additional file 1: Table S3)**. CAL-33 has a missense mutation in the *PIK3CA* gene (g.179234297A>G in GRCh38, p.His1047Arg). BEstimate gRNA design enabled the identification of a new gRNA due to the generation of a novel PAM with the mutated G nucleotide (TGAAACAAATGAATGATGCACGT, GRCh38 genomic location: 179234276-179234298 on chromosome 3) and editable sites (GRCh38 genomic location: 179234278, 179234279, 179234280, 179234282, 179234283, 179234284, on chromosome 3, details in **Additional file 1: Table S3**). The library with the reference sequence prevents the manipulation of the 179234282 genomic location, which allows ABE to generate a missense mutation, p.Gln1042Arg, on the PI3K-PI4K catalytic domain in the *PIK3CA*-encoded p110α subunit.

In summary, BEstimate can design base editor gRNAs for both reference and altered gene sequences, enabling flexibility in both base editor properties and gene sequences, and we illustrate these features by designing a cancer-relevant gRNA library.

### Case study II: Functional annotation of a *MYC* base editor screen

BEstimate utilises the *Ensembl* REST-API to use *Ensembl* Variant Effect Predictor (VEP) to collect the effects of variants (39,45). To illustrate the *in silico* annotation functionality, we obtained data from *MYC* ABE and CBE base editor fitness screens in the colon cancer cell line HT-29 and annotated the gRNA effects (13) **(Additional file 1: Table S4)**. *MYC* is an oncogene and is an essential gene for HT-29 cells. We linked the potential consequences from VEP to the measured viability changes. As expected, while synonymous mutations did not cause a reduction in cell viability, highly depleted gRNAs were predicted to cause missense and nonsense mutations **(Fig. 2c)**. While 72% of the gRNAs were predicted to generate at least one alteration reported in the *Catalogue Of Somatic Mutations In Cancer* (*COSMIC*) (46), 43 gRNAs were predicted to alter a post translational modification sites (acetylation sites at 158, 163, 172, 332 and 338, and phosphorylation sites at 21, 23, 73, 77, 86, 176 and 308) **(Fig. 2c)**. Additionally, we identified 10 gRNAs that generate mutations overlapping human genetic variation sites **(Fig. 2c)**, retrieved from the *Genome Aggregation Database* (*gnomAD*) (47). Naturally occurring variations can lead to potentially different effects across different populations; thus, this result indicates the importance of cautious interpretations, especially in clinical applications.

MYC is a transcription factor and binds DNA through heterodimerisation with MAX (48–50). Blocking MYC-MAX protein interactions negatively affects cancer cell growth (51–54). Concordantly, BEstimate predicts that the highly depleted gRNA (GACGGACAGGATGTATGCTGTGG for ABE, GRCh38 genomic location: 127740834-127740856 on chromosome 8, p.Ser420Pro, **Additional file 1: Table S4**) results in a missense mutation, adding proline, on the helix-loop-helix motif (Log2 Fold Change-LFC = −4.367) at the interface between MYC and MAX **(Fig. 2c)**. Moreover, BEstimate further collects predicted variant consequences from various tools such as Sorting Intolerant From Tolerant (SIFT) (55), Polymorphism Phenotyping (PolyPhen) (56) and Combined Annotation Dependent Depletion (CADD) (57) via the *Ensembl* VEP API. We facilitated categorical SIFT prediction of *MYC* mutagenesis and found that tolerant and deleterious SIFT categories broadly differentiated the differences in *MYC* depletion (two-sided Mann-Whitney U Test, p-value=2e-10, **Fig. 2d**), with some exceptions, highlighting the importance of experimental screens. In summary, BEstimate integrates functional and structural interpretation to elucidate mechanisms of variant effects.

### Case study III: Identifying gRNAs to correct a sickle cell disease-causing variant

In addition to the functional analysis of alterations on the wild-type genomic sequences, using base editors to revert disease-associated pathogenic mutations is clinically promising (58–60). There are examples of clinical stage studies in which disease-causing alterations can be corrected with base editing, such as in sickle cell disease, β-thalassemia (15,16) and Alpha-1 Antitrypsin Deficiency (17). In addition to enabling gRNA design to target SNVs, BEstimate can identify gRNAs predicted to revert disease-associated alterations. As an example, we retrieved the β-globin gene (*HBB*) missense mutation (g.5227002A>T in GRCh38, p.Glu7Val) causing sickle cell disease (61), and identified an NG PAM-specific gRNA for ABE (ACTTCTCCACAGGAGTCAGATGC, GRCh38 genomic location: 5226994-5227016 on chromosome 11). This gRNA enables the mutant codon GTG (Valine) to be changed to GCG (Alanine) and generates the non-sickling variant, Makassar, haemoglobin **(Additional file 4: Fig. S1)** (15,62). This capability could be extended to other monogenic diseases that could be amenable to therapeutic genome editing.

## Discussion

The broad applications of base editors require robust gRNA library design tools with flexible utility and comprehensive annotations for different experimental purposes. Disadvantages of current tools are their limitations on user parameters and limited annotation of the effects of programmed variants. In contrast, BEstimate offers full customisation, accommodating all current and future base editor enzymes and Cas9 proteins while incorporating off-target annotations and extensive functional, structural, and clinical annotations of predicted alterations on genes and proteins. Generating non-reference sequences with the provided list of alterations, BEstimate can generate gRNAs targeting mutant genes, including disease-associated variants and SNPs in genomes of individuals from diverse ancestries. By using APIs, BEstimate ensures up-to-date real-time data retrieval, eliminating the need for manual sequence collection and minimising potential errors. In addition to designing libraries for reference gene sequences, BEstimate can also identify gRNAs for mutant gene sequences and design gRNAs to revert pathogenic variants.

To highlight the functionality of BEstimate, we designed gRNA libraries using various PAM sequences and base editors and explored their coverages and predicted effects. Due to the inherent constraints of base editors, requiring specific PAM sequences and at least one editable nucleotide within a defined activity window, the editable sites across the genome are limited. Similarly, due to the redundancy of the genetic code, alterations might not end up as nonsynonymous alterations. Expanding base editing capabilities with Cas9 variants with less stringent PAM specificity and different base editors targeting different nucleotides can improve the coverage across both nucleotide and amino acid sequences. BEstimate allows users to easily generate gRNA libraries tailored to different editing constraints, such as PAM sequence flexibility, editable window size, and editor type, enhancing coverage at both the nucleotide and amino acid levels.

Cellular models used in base editing experiments might intrinsically harbour alterations in their sequences that affect the presence or absence of gRNA-editable sites. Additionally, these alterations can result in disease conditions, and base editors can be leveraged to manipulate changes into a non-disease genotype. The SNP-aware functionality of BEstimate enables the design of gRNAs not only for reference genomes but also for mutated sequences, making it highly suitable for disease models and personalised editing strategies. BEstimate can identify gRNAs either introducing functional mutations for screening purposes or designing gRNAs capable of reverting disease-causing variants. The flexibility, precision, and extensive annotation make BEstimate a powerful tool for diverse applications, including functional genomics and therapeutic genome editing.

Base editors induce precise genomic changes, which might have significant consequences. Annotating the functional, structural and clinical implications of potential edits helps to interpret the base editing experiment results and to link genomic changes to phenotypic outcomes. The *MYC* screen analysis in this study demonstrated the utility of BEstimate by identifying gRNAs predicted to induce deleterious mutations, impact post-translational modifications, or affect functional domains and protein interaction interfaces. While *in silico* annotation can reveal the functional consequences of experiments, it can also aid the design of base editing libraries before starting the experiment, focusing on edits with specific predicted functional outcomes. This comprehensive annotation property distinguishes BEstimate from previous tools by providing a more informative approach to interpreting results and strategically designing targeted edits.

BEstimate generates gRNA libraries by giving equal probabilities for all positions within the activity window and treating all identified positions as editable. However, previous studies indicated that editing efficiency varies depending on sequence context (63,64), Cas9 protein variant and positional preferences (31,63). Therefore, generating the intended alteration with high efficiency depends on the model, base editor, and gRNA sequence, and necessitates experimental optimisation (63). Additionally, BEstimate currently retrieves gene sequences exclusively from the human genome within the *Ensembl* database. Future enhancements could include expanding the compatibility of BEstimate with additional genomes and non-coding regions and coupling it with base editing efficiency prediction algorithms (63,64). Moreover, tools enabling the integration of three-dimensional locations of the editable positions on protein structure will further extend the application for drug design and protein engineering (65,66).

In summary, BEstimate provides a systematic approach for flexible design and annotation of gRNA libraries for base editing mutagenesis screens and therapeutic applications.

## Conclusion

In this study, we present BEstimate, a versatile Python module that provides a comprehensive approach for designing and annotating base editor gRNA libraries, addressing key limitations of existing tools. Full customisation on defining base editor and gene sequences, integration of *Ensembl* and *Uniprot* APIs for real-time data retrieval, dual functionality on reference and altered sequences, and extensive annotations make BEstimate a valuable tool for diverse research applications.

## Methods

### Overview of BEstimate workflow

BEstimate is a Python-based command line tool (see all inputs in **Additional file 1: Table S5**) to find and annotate Base Editor gRNAs.

**gRNA Template Formation:** For any type of Base Editor, BEstimate can design a representative template via regular expression using user input as the target and edit nucleotides, protospacer length, PAM specificity and location according to the protospacer, and activity window indices on the protospacer **(Fig. 1)**.

**Sequence Preparation:** BEstimate can retrieve the sequence of the interested Hugo Symbol through *Ensembl* REST-API (45) from the user-defined *Ensembl* genome assembly (40) **(Fig. 1)**. In the case where the user may want to integrate the interested one or several SNPs, BEstimate can alter the gene sequences. The format of the SNP should be in Human Genome Variation Society (HGVS) format (67).

**gRNA Identification:** By searching the sequence with the template, BEstimate firstly finds all possible gRNA target sites from both directions. It then uses the Base Editor specification from the user to select the gRNA having at least one editable nucleotide inside the defined activity window **(Fig. 1)**. BEstimate afterwards collects *Ensembl* Transcript, Exon and Protein IDs if gRNAs or editable nucleotides are on them as well as labels the gRNAs if they have a poly-T region.

**gRNA Annotation:** The gRNAs can be annotated functionally, clinically and structurally **(Fig. 1)**. BEstimate utilises *Ensembl* REST-API to use *Ensembl* and *Ensembl* VEP (68) data to find genomic, transcriptomic and proteomic positions of the potential edits. If an *Ensembl* Transcript ID is not provided, BEstimate uses the canonical transcript selected with Matched Annotation from *NCBI* and EMBL-EBI (*MANE*) (69) with a *RefSeq* (70) match. To retrieve the effects of the potential edits with the gRNAs, BEstimate first prepares the Sequence Variant Nomenclature as HGVS (67) of all single and multiple edits. For single edits, substitution versions of HGVSs are prepared; on the other hand, for multiple edits, BEstimate uses indel nomenclature by deleting the reference and inserting the altered activity window sites of the gRNAs. Then, it utilises HGVS symbols as inputs to VEP API to collect information on regulatory regions, DNA motifs and bound transcription factors, mutational consequences, locations of the alterations on genome, cDNA and protein sequences, corresponding *Uniprot* accession, predicted and clinical effects of the alterations and corresponding *Clinvar* (71) and *COSMIC* (72) IDs, and lastly ancestral allele for the given variants from *gnomAD* (47). For the protein level, BEstimate also collects positions of domains and post-translational modification sites (phosphorylation, ubiquitination, methylation and acetylation) via *Uniprot* API (41,42) after aligning the *Ensembl* Protein and *Uniprot* indices. If *Ensembl* API is not providing the index mapping, BEstimate manually maps those two sequences (Python BioPython package (73)) and provides the alignment table as a Comma-Separated Values (CSV) file. These indices are used to label the alteration on protein-protein interaction interface sites, which have been collected from the *Interactome Insider* database (74). BEstimate, thus, maps the positions of corresponding proteins with the genome positions. It therefore merges nucleotide alterations onto protein consequences.

**Off Target Identification:** BEstimate facilitates the CRISPR-Analyser tool from the Wellcome Sanger Institute Cellular Informatics Group (75). First, the chromosomes of the human genome are downloaded from *Ensembl* and indexed using the provided PAM sequence. Then, for each gRNA found by BEstimate, the CRISPR Analyser identifies off-target information **(Fig. 1)**.

### Designing base editor libraries for 50 frequently mutated genes across pancancer

The *cBioPortal* (44) was used to retrieve 50 frequently mutated genes, whose nucleotide length is under 500,000 base pairs, from a pan-cancer study composed of 25,000 patients (43) **(Additional file 1: Table S2)**. Two NG- and NGG-PAM-specific gRNA libraries were designed, and *in silico* annotation was applied for 50 genes with the default parameters except activity window, which was arranged as 3-9, respectively.

### Designing a base editor library for the CAL-51 cell line-specific *PIK3CA* gene

A nonsynonymous single-nucleotide alteration in *PIK3CA* of a head and neck cancer cell line, CAL-33, was collected from the Cell Model Passport (76). A mutation file with its HGVS symbol (3:g.179234297A>G) was created. Then, BEstimate was used to generate the *PIK3CA* library specific to the CAL-33 cancer cell line with default settings except for the PAM sequence (NG) and activity window (3–9) and annotate the library with functional, structural, clinical, and off-target information.

### Identification of gRNAs correcting sickle cell disease-associated *HBB* mutation

The genomic location of the missense mutation causing Sickle cell disease (61) was retrieved from *dbSNP* (rs334) (77). Collected mutation was used to generate mutation files with HGVS symbols from genomic position and nucleotide changes. BEstimate was run for the *HBB* gene for NG-specific adenine base editor (activity window: 3-9) with the prepared mutation file. The gRNAs annotated with ‘guide_change_mutation’ were collected as sequences targeting and manipulating the mutated nucleotide.

### *In silico* annotation of *MYC* base editing screen

The results of *MYC* ABE and CBE screenings were collected from a previous study within the group (13). The *in silico* annotations of the gRNAs were identified with BEstimate using VEP functionality leveraging *Ensembl* VEP REST API (39,45).

## Abbreviations

gRNA: guide RNA
CRISPR: Clustered Regularly Interspaced Short Palindromic Repeats
HDR: Homology-Directed Repair
Indel: insertion and deletion
CBE: Cytosine Base Editor
ABE: Adenine Base Editor
CGBE: Glycosylase Base Editor
Regex: Regular Expression
API: Application Programming Interface
REST: Representational State Transfer
LFC: Log2 Fold Change
gnomAD: Genome Aggregation Database
COSMIC: Catalogue of Somatic Mutations in Cancer
PPI: Protein-Protein Interaction
SIFT: Sorting Intolerant From Tolerant
SNV: Single Nucleotide Variant
VEP: Variant Effect Predictor
PolyPhen: Polymorphism Phenotyping
CADD: Combined Annotation Dependent Depletion
HBB: β-globin gene
SNP: Single Nucleotide Polymorphism
HGVS: Human Genome Variation Society

## Additional Files

Additional file 1: Excel document collecting supplementary tables.

Table S1: Comparison with the previously designed tools.
Table S2: The 50 selected frequently altered genes and their lengths.
Table S3: gRNA library for CAL-51-specific *PIK3CA* gene.
Table S4: *In silico* annotation of the *MYC* screen. Table S5: BEstimate inputs.
Additional file 2: gRNA library for 50 frequently mutated genes, retrieved from a *cBioPortal* pan-cancer study, for all coding and non-coding regions.
Additional file 3: gRNA library for 50 frequently mutated genes, retrieved from a *cBioPortal* pan-cancer study, for all coding regions with protein level annotations.
Additional file 4: PDF document containing the supplementary figure.

Fig. S1: Conversion of sickle cell variant into Makassar, non-sickling variant.

## Declarations

### Ethics approval and consent to participate

Not applicable

### Consent for publication

Not applicable

### Availability of data and materials

The source code is freely available at https://github.com/CansuDincer/BEstimate under the AGPLv3 licence.

Project name: BEstimate

Project home page: https://github.com/CansuDincer/BEstimate

Archived version: BEstimate 1.0.0

Operating system: Linux x86_64

Programming language: Python

License: AGPLv3

The Jupyter Notebooks used to generate the results are archived on Zenodo and can be accessed via the following DOI: https://doi.org/10.5281/zenodo.15364750. All data generated or analysed during this study are included in this published article and its additional files, which are available in the figshare repository with the identifier https://doi.org/10.6084/m9.figshare.28582844.v1 for additional files and https://doi.org/10.6084/m9.figshare.28582916.v1 for the additional figure.

### Authors’ contributions

C.D., M.A.C., and M.J.G. conceptualised the study. C.D. developed the BEstimate Python package and performed the computational analysis and visualisation. B.F. and C.D. adapted the off-target identification pipeline. C.D., M.A.C., and M.J.G. wrote, reviewed, and edited the manuscript. M.A.C. and M.J.G. supervised the study. M.J.G. funded the research. All authors read and approved the final manuscript.

## Acknowledgements

We thank the Garnett laboratory and the Cellular Informatics Group at the Wellcome Sanger Institute for their assistance and Katrina McCarten and Gabriele Picco for their helpful discussions on the design and results of BEstimate.

